# Efficient plasmid-based rescue of T7 RNA polymerase-driven calicivirus reverse genetics systems in mammalian cells using vaccinia virus RNA capping enzymes

**DOI:** 10.64898/2026.03.19.712921

**Authors:** Frazer J.T. Buchanan, Markella Loi, Charlotte Chim, ShuXian Zhou, Rebekah Penrice-Randal, Leandro X. Neves, Maximillian Erdmann, Edward Emmott

**Author notes:** These authors contributed equally (Markella Loi, Charlotte Chim, ShuXian Zhou).

## Abstract

The caliciviruses include important human and animal pathogens such as norovirus, sapovirus and feline calicivirus. Viral reverse genetics is performed to understand the fundamental biology of these viruses, as well as a potential route to generate live-attenuated vaccines. Calicivirus reverse genetics systems have typically relied on either on the production of *in vitro-*transcribed RNA or plasmid-based rescue either from a mammalian promoter, or through supplementing with helper enzymes through means of a helper virus. Here, we present a novel system integrating vaccinia capping enzymes D1R and D12L encoded on plasmids as part of a system for Murine Norovirus (MNV) reverse genetics. Addition of D1R, D12L and T7 RNA polymerase-expressing plasmids increases the viral titres of rescued MNV in both BSR-T7 cells and transgenic BSR-T7^CD300LF^ cells, and viral polyprotein abundance. When the murine norovirus receptor is expressed in BSR-T7^CD300LF^cells, viral titres increased 100-1000-fold compared over standard BSR-T7 cells. This system offers a robust, high-throughput means of assessing viral mutants.

## Introduction

Human norovirus is far more than a transient “stomach bug”, manifesting as the leading cause of acute gastroenteritis worldwide, thus representing a persistent challenge to healthcare systems and a major concern to public health (Winder et al., 2022).Transmission occurs mainly through the faecal–oral route and is facilitated by the virus’ high infectivity and rapid transmission rates (Winder et al., 2022). These characteristics enable norovirus to spread across communities and settings such as care homes and schools, where outbreaks can place a significant strain to health care resources.

Recent epidemiological surveillance data in the United Kingdom suggests shifting patterns in norovirus activity, with notable increases in confirmed cases since 2023 and continued circulation of the virus in following years, particularly during the winter months (UKHSA, 2026). Such trends emphasise norovirus’ everchanging dynamics of transmission and serve as a reminder of the significance of sustained monitoring and research efforts. With no licensed vaccines or antivirals, stricter hygiene practices serve as the main preventative measures against norovirus (Omatola et al., 2024). Additionally, the continuous emergence of new variants like GII.17 further complicate and delay the development of effective therapeutics (Doh et al., 2025). Progress towards these goals relies not only on epidemiological surveillance, but also on detailed understanding of the molecular mechanisms that modulatenorovirus replication and theconserved biological events within its viral family, the *Caliciviridae*.

Caliciviruses are characterised by linear single-stranded, positive-sense RNA genomes of 7.4-8.3 kb, and depending on their genus, typically possess either two or three opening reading frames (ORFs) (Téllez et al., 2024; Vinjé et al., 2019). The non-structural (NS) proteins 1-7 (NS1-7) are encoded as a large polyprotein from ORF1 (∼7.4 kb) beginning at the 5’ end of the genome. Genes encoding the structural proteins are located at the 3’ end of the genome, and encoding major structural protein 1 (VP1) and minor capsid protein (VP2),. Processing of the polyprotein is mediated by both the viral 3CL protease NS6 and host caspases; this process is essential to allow completion of the replication cycle (Emmott et al., 2019, 2015) (Figure 1A). Viral protein genome-linked (VPg), once liberated from the polyprotein, serves as a primer for the norovirus genomic and subgenomic RNA, as well as a cap substitute for translation of viral proteins (Goodfellow et al., 2005; Olspert et al., 2016; Royall and Locker, 2016).

**Figure 1.**
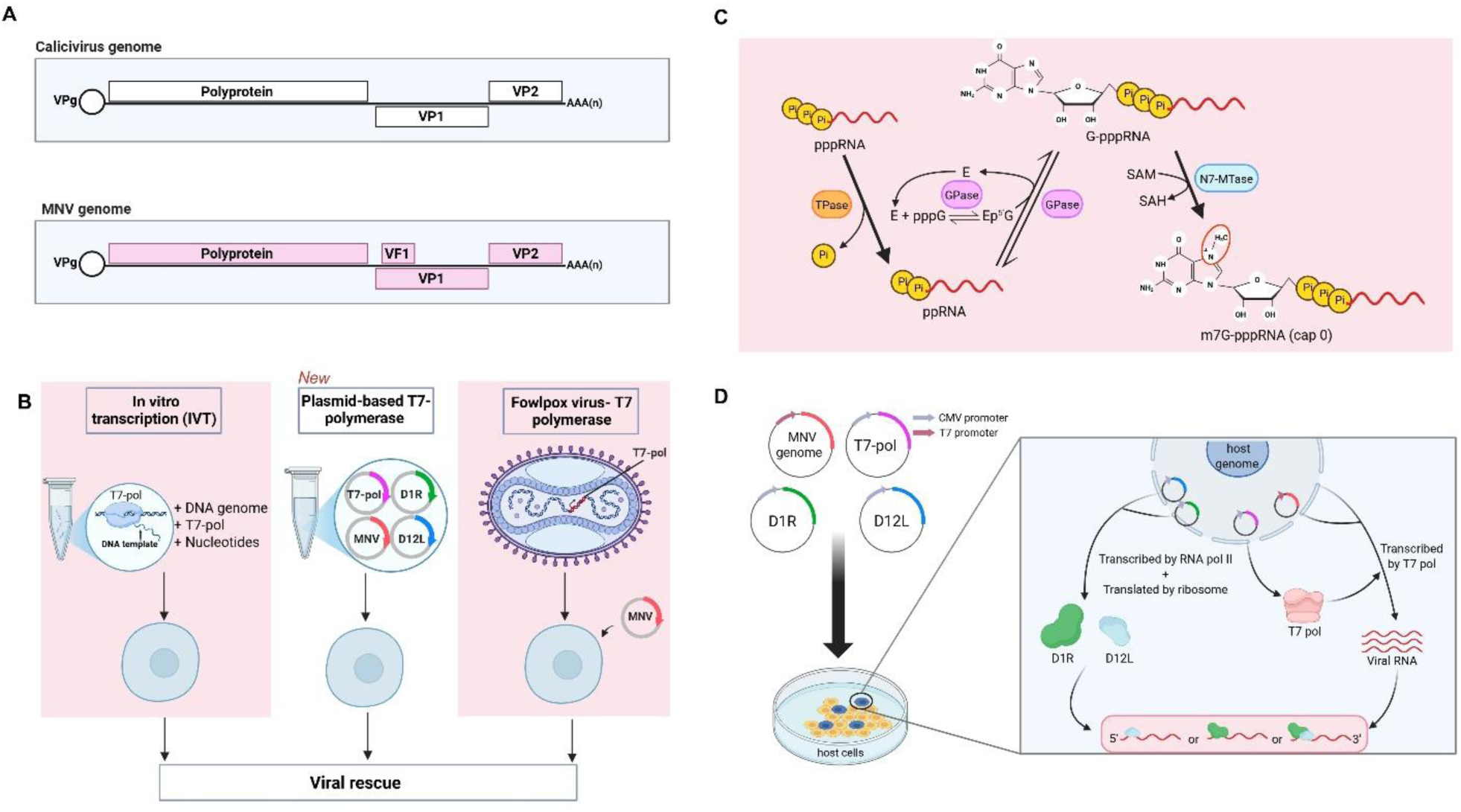
Summary of T7-based systems for Calicivirus reverse genetics. **A)** Schematic outlining a generic calicivirus murine norovirus (MNV) genome, including polyprotein, virulence factor 1 (VF1), viral protein 1 (VP1) and viral protein 2 (VP2), viral protein genome-linked (VPg). **B)** Outline of current T7-based reverse genetics systems (pink panels) including IVT and Fowlpox-T7 based rescue. The middle panel outlines a reverse genetics system incorporating vaccinia virus capping proteins D1R and D12L. **C)** Canonical RNA 5’ capping: triphosphorylated RNA (pppRNA) is converted to dephosphorylated RNA (ppRNA) by an RNA 5’ triphosphatase (TPase). Guanylytransferase (GTase) forms a covalent enzyme intermediate (E-pG) and transfers GMP to the 5’ end of ppRNA, generating GpppRNA. Cap structure is then methylated at the N7 position of the guanine by methyltransferase (N7 - MTase) using S-adenosylmethionine (SAM) as the methyl donor, producing m^7^GpppRNA. **D)** A combination of genome and helper plasmids (D1R, D12L & T7-pol) would allow non-canonical capping of T7 RNA transcripts produced intracellularly.

Despite their long-standing clinical and epidemiological significance, progress in understanding the molecular biology of caliciviruses has been comparatively slow (Téllez et al., 2024). A major limitation in calicivirus research has been the difficulty of propagating these viruses in cell culture and thelack of animal models (Desselberger, 2019; Karst, 2010). Therefore, much of what is known about calicivirus biology has been derived from a small number of experimentally tractable model viruses (Goodfellow and Taube, 2016). Chief among these are feline calicivirus (FCV) and murine norovirus (MNV) (Peñaflor-Téllez et al., 2022). Especially the latter has emerged as the most widely used surrogate for human norovirus, a case attributed to MNV’s efficient replication in tissue culture and the availability of a well-established small animal model (Karst, 2010).

Within these model systems, the scope of experimental analysis is shaped by the available molecular tools. For positive-sense RNA viruses, and particularly caliciviruses, these systems have traditionally relied on the delivery of *in vitro-*transcribed (IVT) genomic RNA (Hoffmann et al., 2024). Full-length viral RNA is synthesised *in vitro,* typically using bacteriophage RNA polymerases, and transfected directly into cells that can initiate infection (Lenk et al., 2024). IVT-based reverse genetics has been instrumental in establishing foundational knowledge and remains a reliable method for generating infectious virus (Lee et al., 2023; Mu and Hur, 2021). They are often considered the gold standard system for viral reverse genetics; however, they require additional steps such as synthesis compared to solely plasmid-based systems. The requirement for repeated synthesis of high-quality RNA, coupled with its susceptibility to degradation and the need for careful quality control, can limit experimental throughput (Lee et al., 2023; Lenk et al., 2024). These constraints become particularly restrictive when large numbers of viral variants must be generated, making this method poorly suited to scalable or iterative experimental designs.

Reverse genetics systems have been pivotal to advancing virological research, allowing viral rescue from purified DNA or RNA, and thus enabling the artificial engineering of viral genes (Chen et al., 2022; Zhang et al., 2025). In calicivirus research, this approach was first successfully demonstrated for FCV in 1995, confirming recovery of the virus from a genetically engineered mutant (Sosnovtsev and Green, 1995). Nonetheless, the inability of the obtained viral transcripts to consistently and successfully trigger virus replication and recovery has led to the limited use of reverse genetics systems for caliciviruses – with the field remaining at its infancy (Álvarez et al., 2024; Goodfellow and Taube, 2016).

An alternative approach for calicivirus propagation, specifically MNV- uses recombinant fowlpox virus (FWVP) expressing T7 RNA polymerase which permits recovering of virus from DNA based constructs (Chaudhry et al., 2007). Chaudhry et al., established that MNV reverse genetics posed distinct challenges compared with other caliciviruses and that vaccinia virus-based systems were incompatible with efficient MNV replication, whereas fowlpox virus provided a permissive context. Although this system represents a major advance, its reliance on helper virus infection and propagation in primary chicken embryo fibroblasts introduces technical limitations, that restrict accessibility as well as scalability. These considerations highlight the broader need for reverse genetics systems that retain biological fidelity while addressing complexity.

Plasmid-based reverse genetics strategies could offer an alternative for initiating viral replication by allowing intracellular transcription of viral genomes from DNA templates. Here, we introduce a plasmid-based reverse genetics system for MNV that combines a T7-driven viral genomewith helper plasmids encoding thevaccinia virus mRNA capping enzyme subunit D1R and its cofactor D12L (Ramanathan et al., 2016; Tate et al., 2016) (Figure 1). This system is designed to enhance the production of translation-competent viral RNA while avoiding the need for IVT or helper virus propagation. All components are delivered as plasmid DNA, allowing rapid exchange of viral genome constructs and facilitating experimental workflows that require repeated or parallel manipulation. This feature is particularly relevant for studies that require comparison of multiple variants and may encourage the adaptation of reverse genetics strategies to other caliciviruses that remain difficult to manipulate.

Beyond its immediate application to MNV, this strategy could provide the framework for a more accessible and scalable platform for routine viral reverse genetics, complemented by proteomic profiling and may inform the design of reverse genetics systems for other caliciviruses.

## Materials & Methods

### Cell culture

Cells were tested at monthly intervals for mycoplasma and confirmed negative for mycoplasma infection. BV-2 cells were a gift from Prof. Ian Goodfellow, University of Cambridge and BSR-T7 herein referred to as BSR-T7, Professor Julian Hiscox, University of Liverpool. Cells were cultured in DMEM (Sigma-Aldrich) supplemented with 10 % Foetal Bovine Serum (FBS) (Sigma-Aldrich) and 50 U ml^-1^ penicillin and 50 µg ml^-1^ streptomycin. All cells were maintained at 37 °C with 5 % CO_2_. BV-2 cells were used in this study for MNV TCID50s. BSR-T7 cells were used in this study for viral reverse genetics. CD300LF-positive BSR-T7 cells (BSR-T7^CD300LF^) were maintained in 2.5 µg ml^-1^ of puromycin (Gibco).

### Plasmids

The vaccinia capping plasmids, pCAG-D1R, pCAG-D12L were gifts from Takeshi Kobayashi (Addgene plasmids #89160 and #8916 (Kanai et al., 2017). The T7opt in pCAGGS plasmid was a gift from Benhur Lee (Addgene plasmid #65974) (Yun et al., 2015). The MNV reverse genetics system pT7 MNV 3’RZ was a gift from Ian Goodfellow, University of Cambridge (Chaudhry et al., 2007; Yunus et al., 2010). The pGenLenti-murine-CD300LF plasmid for lentiviral-based expression of murine CD300LF was synthesised by Genscript and is available from Addgene (Addgene plasmid #235668).

### Generation of CD300LF expressing BSR-T7 cells(BSR-T7^CD300LF^)

BSR-T7 cells were seeded (3×10^5^ cells/ well) into 6 well plates 24 h prior to transduction with Lentiviral vector (pGenLenti-CD300LF) (Genscript). Cells were treated for 30 mins with 8 µg ml^-1^ of polybrene prior to transduction of lentivirus at an MOI of 1. Cells were then put under puromycin selection at a variable concentration 2-4 µg ml^-1^ and cell death monitored daily. CD300LF expression was maintained by culturing cells in 2.5 µg ml^-1^ puromycin.

### Viral rescue

MNV pT7 MNV 3’RZ plasmids were transfected into BSR-T7 cells, alongside pCAG-D1R, pCAG-D12L, T7opt at a 1:1:1:1 ratio. BSR-T7 cells were reverse transfected using lipofectamine 2000 (Invitrogen) and a total of 2 µg of plasmid DNA. 6 h following transfection cell culture media was changed. Plates were then incubated for a total of 72 h Culture plates containing the transfected cells then underwent a freeze-thaw cycle at –80 °C.

### Viruses

Full-length MNV clone based on the NC_008311.1 Norovirus GV, sequence under the control of a T7 polymerase promoter (Chaudhry et al., 2007). All MNV infections were conducted in BV-2 cells.

### TCID_50_

Quantification of MNV particles was conducted via tissue culture infectious dose 50 assay in BV-2 cells. Virus was harvested by freezing cells and media at -80 °C, allowing lysis through a freeze-thaw cycle. Lysates were then serially diluted 10-fold in cell culture media and added to BV-2 cells in a 96-well plate and incubated for 120 h at 37 °C with 5 % CO_2_. The TCID_50_ was calculated by determining the dilution factor required to show 50 % CPE (Hwang et al., 2014). TCID_50_ results were visualised in RStudio (version 2025.09.2+418).

### Western blotting & antibodies

Cell were lysed in 1X RIPA lysis buffer (1M Tris-HCl, 5M NaCl, 2.5 % Sodium Deoxycholate, 10 % NP40, 10 % Sodium dodecyl sulphate, ddH_2_O, supplemented with HALT cocktail protease inhibitor 1:200 (Thermo scientific) and lysates were centrifuged for 15 mins at 15000 g, clarified supernatants were retained in separate tubes. 3.5 µl of 4X Lamelli buffer (BioRad) supplemented with β-mercaptoethanol (10%) was added to samples. Proteins were separated on a 17.5 % polyacrylamide SDS gel and subsequently transferred onto PVDF membrane (Immobilon) using a BioRad Trans-Blot turbo transfer system. Membranes were blocked in 10 % milk in Tris-buffered saline solution supplemented with 0.1 % Tween (TBS-T, Sigma-Aldrich). Membranes were then incubated at 4°C overnight with gentle agitation with rabbit anti-VpG antibody (1:1000) (In-Custom Polyclonal generated by Eurogentec). Membranes were washed three times in TBS-T prior to incubation with anti-rabbit HRP-conjugated secondary antibody (1:2000) (Cell signalling). Membranes were washed three times in TBS-T and incubated in Pierce© ECL reagent (Thermo Scientific) prior to imaging on a ChemiDoc MP System (BioRad).

### Proteomics

BSR-T7 cells were harvested and washed twice in pre-warmed Dulbecco’s phosphate buffered saline (DPBS; Sigma Aldrich) then lysed in 100 mM HEPES pH 7.4 containing 1 % IGEPAL CA-630, 1 % sodium dodecyl sulphate and 1x Halt protease inhibitor cocktail (Thermo Scientific) and heated at 95 °C for 5 mins, 900 rpm. Benzonase Nuclease (Sigma Aldrich) was added at 1:200 dilution (v/v) following 15 min incubation at 37 °C, then dithiothreitol (DTT) was added to 4 mM final concentration and samples heated at 60 °C for 10 mins, 900 rpm. The samples were allowed to cool to room temperature and iodoacetamide (IAM) was added to 14 mM final concentration following 30 mins incubation at room temperature (RT) in the dark. The excess of IAM was then quenched with a second addition of 3 mM DTT to samples. Protein concentration was measured using Pierce™ BCA Protein Assay Kit (Thermo Scientific) following the manufacturer’s microplate procedure. 15 µg of each sample was then transferred to a 96-well LoBind plate (Eppendorf) for SP3-based clean-up and digestion (REF: 10.1038/s41596-018-0082-x). In brief, samples’ volumes were normalised to 50 µl with lysis buffer then 5 µl of 30 µg µl^-1^ Sera-Mag SpeedBead magnetic particles (Cytiva; #45152105050250, #65152105050250) added at a 10:1 bead to protein ratio. Protein was precipitated onto beads by adding 130 µl acetonitrile (MeCN; 70 % (v/v) final concentration) following 15 mins incubation, RT, 750 rpm. In a magnetic rack, beads were washed twice in 220 µl of 90 % (v/v) ethanol, then once in 220 µl of MeCN. To each well, 150 µl of 50 mM ammonium bicarbonate containing 3 ng µl^-1^ Trypsin Gold (Promega; 1:30 protease to protein ratio) following a 2 mins sonication in water-bath to disaggregate the beads. Tryptic digestion was carried out at 37 °C for 16h, 1,100 rpm. In the following day, the digestion supernatant was transferred to a fresh 96-well LoBind plate containing 5 µl of 10 % (v/v) TFA per well, approximately 0.3 % (v/v) final concentration, to terminate the digestion.

LC-MS/MS analysis was carried out using an Evosep One (Evosep Biosystems) coupled to a timsTOF HT mass spectrometer (Bruker). Approximately 150 ng of the tryptic digests was loaded into EV2011 Evotip Pure tips (Evosep Biosystems), as per manufacturer’s instructions. The peptide mixtures were resolved using a EV1906 Endurance Column (Evosep Biosystems; ReproSil-Pur C18, 1.9 µm, 15 cm x 150 µm), 40 °C, and the 30 samples per day (SPD) method. The mass spectrometer was equipped with a CaptiveSpray ion source operating a 1600 V and mass spectra (100 to 1700 *m*/*z* window) were acquired in PASEF (Meier et al., 2018) positivemode. TIMS settingsincluded a 100 ms ramp, ion mobility (IM) coefficients (1/K_0_) from 0.6 to 1.6 Vs/cm^2^. Ten PASEF ramps MS/MS were scanned per cycle, and peptide precursors within the polygon filter applied to the *m*/*z* and IM plane were isolated (2 Th for *m*/*z* < 700 and 3 Th for *m*/*z >* 700) for activation by collision-induced dissociation (CID). Collision energy was linearly ramped as a function of IM, starting at 20 eV until 59 eV across 0.6 to 1.6 Vs/cm^2^. Dynamic exclusion was set to 0.4 mins. Injection order was randomised using the sample() function in R.

Spectral data was processed using FragPipe v22 (Yu et al., 2020) using the “LFQ-MBR” workflow but with MBR disabled. Digestion mode was set to semi-specific trypsin (P1 = KR unless P1’ = P) and up to 1 missed cleavage was allowed. Target-decoy (reverse) was used to estimate false-discovery rate (FDR) and results was filtered at 1 % FDR at the PSM, peptide and protein levels. Database search was performed against UniProt’s Golden Hamster (UP000189706, 20395 entries; downloaded 11/12/2024) and Murine Norovirus 1 (UP000109015, 4 entries; downloaded 08/06/2024) reference proteomes. Data analysis was performed in R using ggplot2 package for box plots and coverage map diagrams. Data are available via ProteomeXchange with identifier PXD074707.

### Data and Code Availability

All code for data analysis is available from https://github.com/emmottlab/T7Capping_2024. The mass spectrometry data has been submitted to ProteomeXchange Consortium (<u>ProteomeCentral Data and Tools</u>) via the PRIDE partner repository with the dataset identifier PXD074707. All manuscript figures are available through Figshare (https://doi.org/10.6084/m9.figshare.31812781).

## Results

### Efficient cell-based rescue from T7-driven calicivirus reverse genetics systems requires co-expression of capping enzymes

Calicivirus reverse genetics systems fall into two categories: 1) Those which are driven by the T7 RNA polymerase promoter and are either synthesised as RNA transcripts and capped *in vitro* or through use of Fowlpox-T7 to drive T7 expression in cells, or 2) Those where expression is driven by a mammalian promoter. The BSR-T7 cell line, which expresses T7 RNA polymerase, is commonly used for viral reverse genetics and viral protein expression (Buchholz et al., 1999; Römer-Oberdörfer et al., 1999; Sandoval-Jaime et al., 2015). In the case of mammalian promoter-driven expression, and fowlpox-T7, either nuclear mammalian RNA capping enzymes, or fowlpox capping enzymes would be expressed within infected cells. However, in the absence of viral capping enzymes, T7 transcription would be expected to result in the synthesis of full-length, albeit uncapped mRNA. As RNA capping is well established as crucial for efficient translation of mRNA into protein, we hypothesised that expression of T7 polymerase alone would result in poor translation of viral proteins from a T7 promoter-driven viral reverse genetics system and, consequently low efficiency of virus recovery.

We tested the hypothesis that cell-based rescue of T7-driven caliciviruses could be achieved by plasmid-based overexpression of T7 RNA polymerase and the vaccinia virus D1R and D12L capping enzymes as helper plasmids alongside the viral T7-driven reverse genetics plasmid. Figure 2A shows the impact of overexpression of individual T7 RNA polymerase, D1R and D12L alongside either wild-type or an inactive polymerase mutant calicivirus reverse genetics plasmid encoding murine norovirus. As expected, all conditions using the YGSN plasmid genome, a plasmid conferring a point mutation within the conserved YGDD active site of the RdRp, which prevents viral replication but can serve as a control for translation from input RNA or plasmid did not produce infectious virus with TCID_50_/ml reading <1 log_10_. This was also the case for the mock condition TCID_50_/ml reading <1 log_10_. The most efficient rescue required expression of all 3 helper plasmids (T7 + D1R + D12L), with infectious virus only, produced from cells transfected with the wild-type viral plasmid, with a 2-fold log_10_ increase relative to the standard WT + T7 condition. Notably the combination of WT genome + D1R & D12L also demonstrated an approximately 2-fold log_10_ increase compared to the WT +T7 only combination. Inclusion of the T7 RNA polymerase plasmid levels of infectious virus, suggesting that the endogenous levels of T7 RNA polymerase in BSR-T7 cells are not enough for efficient viral rescue from plasmid launch.

**Figure 2.**
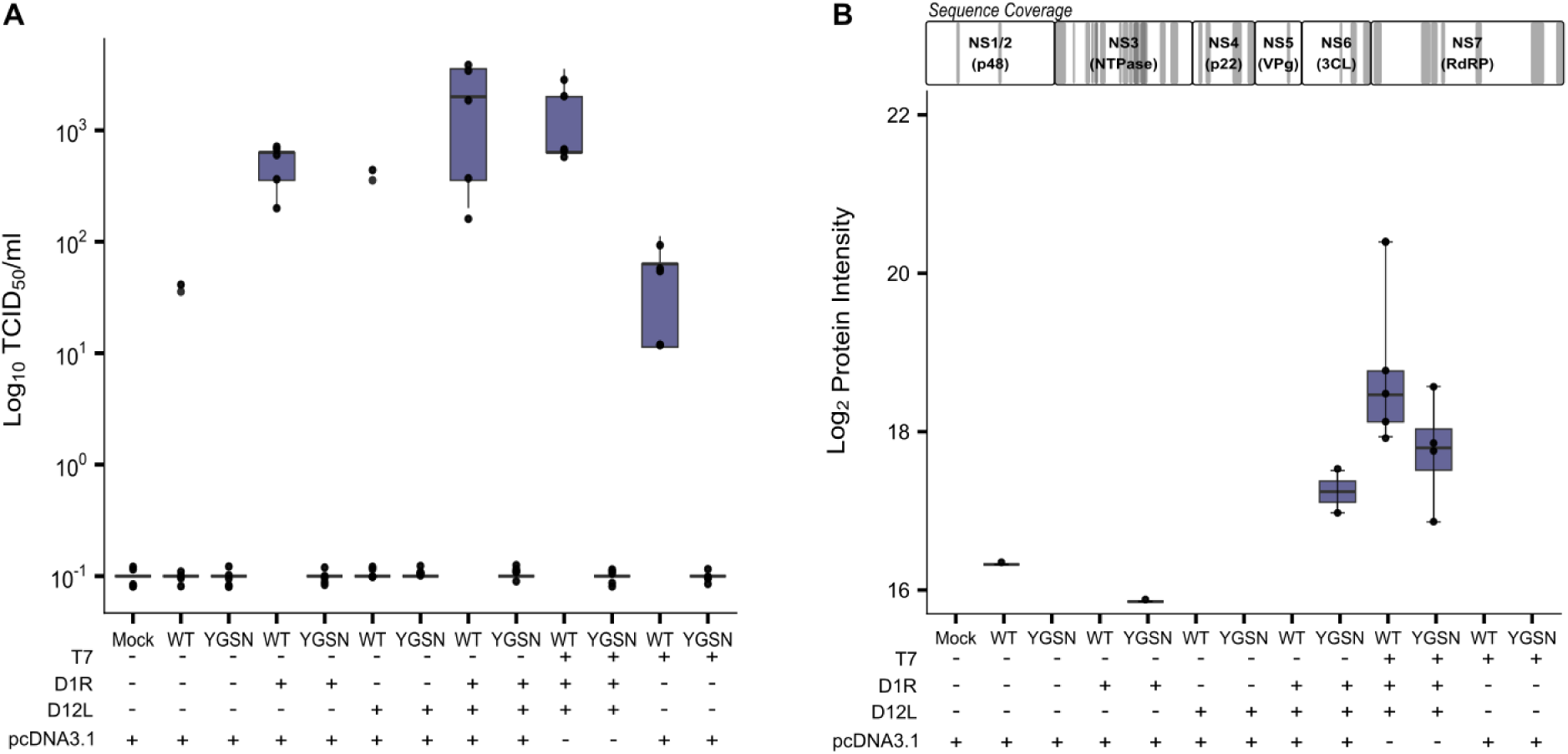
BSR-T7 cells require supplementation with T7 polymerase and both vaccinia capping enzymes for efficient MNV rescue. **A)** TCID_50_ data from BV-2 cells infected with supernatents from transfected BSR-T7 cells, 72h post-transfection with the indicated plasmid combinations. n=4, boxplots show the median (centre line), interquartile range (box; 25th–75th percentiles), and whiskers extending to 1.5× the interquartile range). **B)** LC-MS/MS intensity for protein expression from the viral polyprotein under the indicated conditions. Shaded areas on the diagram highlight the position of peptides identified by LC-MS. The Sequence coverage indicates the peptide origin against viral polyprotein from which peptides were derived (n=5).

We next sought to evaluate the abundance of viral proteins in cells that underwent transfection with different combinations of reverse genetics and helper plasmids by LC-MS/MS. Similarly to Figure 2A, BSR-T7 cells were transfected with a combination of helper plasmids and MNV genome plasmid (WT vs YGSN negative control). Cells were harvested 96 h post-transfection and prepared by SP3 digest for analysis by LC-MS/MS. As expected, the mock condition showed no viral polyprotein expression over the limit of detection. The YGSN combination, including both helper plasmids, resulted in an approximately 18 log_2_ intensity score for polyprotein abundance. This can be explained by the production of translation-competent transcripts in the rescue, which permit protein expression but cannot complete a full replication cycle due to the YGSN defective polymerase mutation. Similarly, the YGSN combination including both helper plasmids and T7 resulted in an approximately 18.5 log_2_ intensity score for polyprotein abundance. Again, this is likely due to the production of viable transcripts which allow input translation of the polyprotein. The WT genome rescue condition including helper plasmids and T7 resulted in an approximately 1.12 log_2_ fold increase in polyprotein abundance compared to the WT genome only. Also notable was that while good sequence coverage of the murine norovirus polyprotein was observed in the LC-MS/MS results, viral proteins expressed from the sub-genomic viral mRNA (VP1, VP2, VF-1) were not observed (Figure 2B).

### Cell-based calicivirus rescue is enhanced using permissive cells

BSR-T7 cells are commonly used for viral reverse genetics based on both their expression of T7 RNA polymerase, but also deletions which impair sensing of plasmid DNA and uncapped RNA (Habjan et al., 2008). BSR-T7 cells are not known to be permissive for MNV infection. However, MNV is able to infect a range of cell lines following expression of its cognate cellular receptor: murine CD300LF (Haga et al., 2016; Orchard et al., 2016). We next sought to understand if transducing BSR-T7 cells to stably express the murine norovirus receptor would yield a more efficient rescue system. Stable expression of murine CD300LF was achieved by lentiviral transduction to deliver CD300LF to the BSR-T7 cells under the control of the CMV promoter, alongside a puromycin-resistance cassette.

To test our hypothesis that expression of murine CD300LF would result in a more efficient virus rescue, we transfected the MNV genome plasmids, along with the T7 RNA polymerase, D1R and D12L helper plasmids into either wild-type BSR-T7 cells or BSR-T7^CD300LF^ cells. As can be seen in Figure 3A, expression of murine CD300LF cells greatly improves the titres obtained at 72 h post-transfection, with a 100-1000-fold increase in average titres, from mid 10^3^TCID_50_/mL to 10^7^TCID_50_/ml.

**Figure 3.**
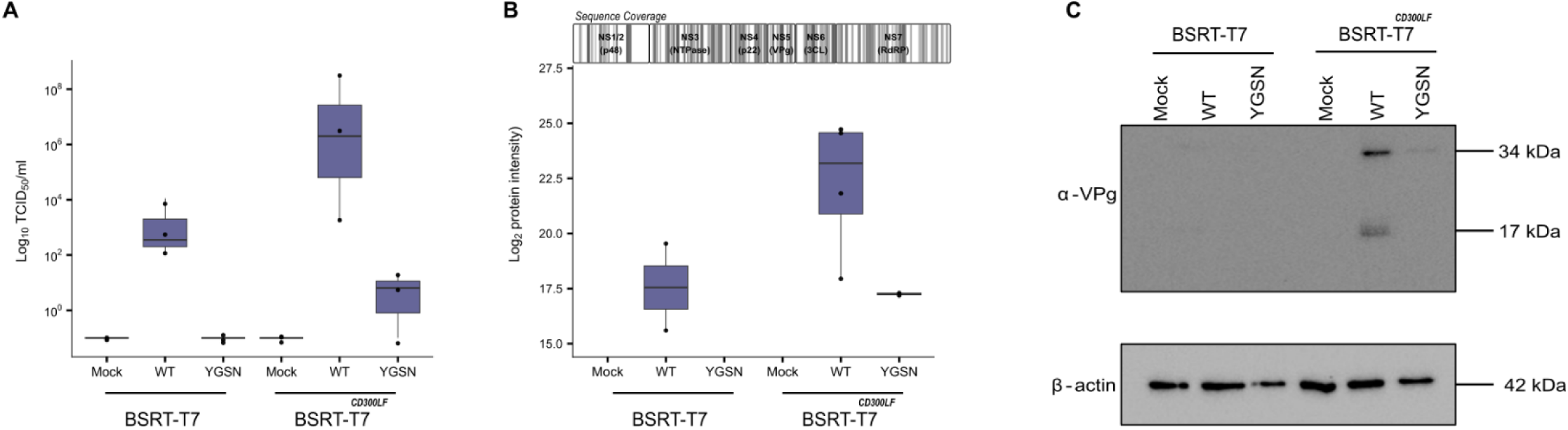
MNV rescue is enhanced by stable expression of CD300LF in BSR-T7 cells. **A)** TCID_50_ data from BV-2 cells infected with supernatents from transfected BSR-T7 or BSR-T7^CD300LF^ cells, 72h post-transfection with the indicated plasmid combinations. n=3, boxplots show the median (centre line), interquartile range (box; 25th–75th percentiles), and whiskers extending to 1.5× the interquartile range). **B)** LC-MS/MS intensity for protein expression from the viral polyprotein under the indicated conditions. Shaded areas on the diagram highlight the position of peptides identified by LC-MS. The Sequence coverage indicates the peptide origin against viral polyprotein from which peptides were derived (n=5). **C)** Western blot analysis of Mock, WT or YGSN-transfected BSR-T7 or BSR-T7^CD300LF^ cells, with anti-VPg antibody.

In agreement with the TCID_50_ data, weobserved a marked increase in VPg expression in BSR-T7 cells expressing CD300LF compared to naïve BSR-T7 cells via western blotting Figure 3C. This is indicated by the stronger bands representing cleaved and Uncleaved VPg (including precursors) detected with anti-VPg antibody.

Given the increase in viral protein abundance when T7 and D1R and D12L areincluded in the rescue setup, we set out to determine polyprotein abundance by LC-MS/MS in both BSR-T7 cells and the BSR-T7^CD300LF^ cells (Figure 3B). We observed an approximately 1.3 log_2_ fold increase in protein intensity for the BSR-T7^CD300LF^ cells compared to the BRST7 cells. This was expected, as the CD300LF expression permits onward replication of virus produced in initially untransfected/infected cells.

## Discussion

Viral reverse genetics systems for positive-sense RNA viruses and more specifically caliciviruses typically rely on the delivery of the viral genome in the form of *in* vitro transcribed (IVT) RNA. An alternative approach involves providing a viral genome encoded within a plasmid under T7 promoter control, coupled with infection of cells using recombinant Fowlpox virus (FPV) expressing T7 polymerase (Chaudhry et al., 2007). Whilst this FPV-based system permits genome expression without direct RNA transfection, it requires propagation of recombinant FPV in primary chicken embryo fibroblasts which can be challenging to access, is labour intensive and can introduce variability due to the primary cell context. IVT RNA-based transfections by contrast remain the gold standard for initiation viral replication and provide precise control over input RNA molecules. However, throughput can be limited in large-scale mutagenic screens and efficiency can be impacted by RNA stability and host cell permissiveness. Reagents for in vitro RNA synthesis can also introduce significant expense to workflows. These limitations in current reverse genetics systems highlight trade-offs among ease of use, scalability, expense, utility and experimental control, underscoring an avenue for alternative rescue strategies.

Plasmid-based approaches for calicivirus reverse genetics employing eukaryotic promoters have been successfully applied previously. Examples include Tet-inducible Pol II-driven murine norovirus reverse genetics (Ward et al., 2007), EF-1-α driven reverse genetics for murine norovirus (Katayama et al., 2014) and feline calicivirus (Oka et al., 2014) and CMV-driven reverse genetics for RHDV (Liu et al., 2008). These systems have the clear advantage of simplicity, using a single or pair of plasmids for transfection, and as they employ mammalian promoters they are capped and exported from the nucleus in line with standard mRNA processing. Disadvantages include often low yields e.g. 10^3^ (Ward et al., 2007), the potential for cryptic splicing, and unclear RNA termini for some mammalian promoters.

Here, we present a novel approach for plasmid-based rescue of MNV, by integrating plasmids encoding the vaccinia virus capping enzyme D1R and cofactor D12L, under T7 expression. Co-transfection of these helper plasmids with a plasmid encoding for MNV genome improved viral recovery in both naïve BSR-T7 cells and those rendered permissive to MNV entry (BSR-T7^CD300LF^) (Figure 3). A YGSN mutant was incorporated to account for the initial noise of input translation from first round genome translation. Both our viral and proteomic assays indicated improved rescue of MNV virions when using our combination plasmid-based assay (Figures 2 & 3). A benefit of the presented rescue system is the ability to rapidly screen MNV mutants; using this plasmid-based system permits the swapping of WT plasmid in the transfection with ease.

Given that caliciviruses utilise VPg to support replication both as a primer and cap substitute for protein synthesis, it is perhaps unsurprising that supplementing RNA capping improved viral rescue (Wei et al., 2008). The importance of VPg or capping for successful viral replication and rescue has long been established precedent (Yunus et al., 2010), highlighting the critical role of 5’ end modifications in positive-sense RNA virus biology. Beyond caliciviruses, supplementation of capping functions has also been shown to improve rescue efficiency in other positive-sense RNA viruses. For example, reovirus reverse genetics systems have been demonstrated to benefit from the presence of a fusion protein (C3P3) encoding for both T7 polymerase and the African Swine Fever Virus (ASFV) capping enzyme NP868R (Eaton et al., 2017). Interestingly, the same NP868R-T7 fusion protein has been adapted for calicivirus reverse genetics, specifically Tulane Virus (TV) (Scribano et al., 2025), demonstrating the versatility of combining RNA polymerase and capping functions to improve viral rescue. These observations suggest that strategies to enhance 5’ functionality whether through VPg or capping enzymes can be broadly applied to improve viral rescue efficiency across multiple positive-sense RNA virus families. By expressing the vaccinia virus capping enzymes D1R and D12L, our system supplements 5’ end capping, fulfilling a key requirement for efficient calicivirus replication and highlighting the broader utility of enhancing cap-like functions in reverse genetics strategies.

A logical next step for the development of this system would be to ensure coordinated and consistent expression of both helper plasmids and the viral genome. Currently, the need to transfect three separate helper plasmids (D1R, D12L & T7) alongside the genome plasmid presents potential issues. Namely, an increase on host cell stress, activation of DNA sensing pathways and competition for translational machinery (Majerciak and Zheng, 2025; Semenova et al., 2019). One strategy to overcome these limitations would be to consider consolidation of all helper plasmids into a single tricistronic plasmid (Peeters and De Leeuw, 2017).

The system described here has the potential to improve viral rescue efficiency of not only MNV but other members of *Caliciviridae*, allowing the opportunity to advance fundamental research. In particular, Human Sapo Virus (hSapo) has been historically difficult to propagate in cell culture due to limited receptor information and inefficient replication in standard host cell lines (Oka et al., 2018; Takagi et al., 2020). The identification of CD36 as an essential host factor for hSapo propagation (Oka et al., 2025) highlights the importance of both cellular context and host factor availability in successful virus propagation. Our system could potentially complement CD36 overexpression, supporting improved sapovirus rescue. More broadly, combining strategies that enhance RNA stability, translation and host factor engagement may facilitate the development of enhanced reverse genetics systems for other caliciviruses.

In summary, we have presented a novel plasmid-based reverse genetics system for MNV which improves viral output in both BSR-T7 and BSR-T7^CD300LF^ cells in culture and we show modification of BSR-T7 cells to express the CD300LF receptor permits high titre rescue from current T7-based reverse genetics plasmids. We also demonstrate the importance of viral genome capping to initiate and improve the efficiency of viral rescue. This system could be expanded into other caliciviruses such as FCV and hSapo, and allows for both plasmid-based and IVT-based viral reverse genetics.

## Acknowledgements

Theauthors thank the other members of the Emmott lab for their support, as well as Dr Philip Brownridge for support with LC-MS/MS instrumentation. EE is funded by an Academy of Medical Sciences Springboard Award (SBF006\1008) supported by the British Heart Foundation, Diabetes UK, the Global Challenges Research Fund, the Government Department for Business, Energy and Industrial Strategy and the Wellcome Trust. The Medical Research Council [MR/X000885/1], BBSRC [BB/W019744/1] and a Wellcome Trust Career Development Award [227831/Z/23/Z]. Dr Christine Tait-Burkard (Roslin Institute) is thanked for her advice on vaccinia plasmids.

## Author contributions

**FJTB, ML, CC, SZ, RPR & ME** prepared samples and conducted data analysis. **FJTB, RPR, ME** and **EE** provided supervision. **FJTB, ML** and **EE** led writing of manuscript. All authors helped finalise the manuscript and was approved by all. **LXN** and **FJTB** led mass spectrometry sample preparation, acquisition and analysis. **EE** conceived and provided funding for the project.

## References

Álvarez, Á.L., Arboleya, A., Abade Dos Santos, F.A., García-Manso, A., Nicieza, I., Dalton, K.P., Parra, F., Martín-Alonso, J.M., 2024. Highs and Lows in Calicivirus Reverse Genetics. Viruses 16, 866. 10.3390/v16060866

Buchholz, U.J., Finke, S., Conzelmann, K.-K., 1999. Generation of Bovine Respiratory Syncytial Virus (BRSV) from cDNA: BRSV NS2 Is Not Essential for Virus Replication in Tissue Culture, and the Human RSV Leader Region Acts as a Functional BRSV Genome Promoter. J Virol 73, 251–259. 10.1128/JVI.73.1.251-259.1999

Chaudhry, Y., Skinner, M.A., Goodfellow, I.G., 2007. Recovery of genetically defined murine norovirus in tissue culture by using a fowlpox virus expressing T7 RNA polymerase. Journal of General Virology 88, 2091–2100. 10.1099/vir.0.82940-0

Chen, H., Liu, H., Peng, X., 2022. Reverse genetics in virology: A double edged sword. Biosafety and Health 4, 303–313. 10.1016/j.bsheal.2022.08.001

Desselberger, U., 2019. Caliciviridae Other Than Noroviruses. Viruses 11, 286. 10.3390/v11030286

Doh, H., Lee, C., Kim, N.Y., Park, Y.-Y., Kim, E., Choi, C., Eyun, S., 2025. Genomic diversity and comparative phylogenomic analysis of genus Norovirus. Sci Rep 15, 5412. 10.1038/s41598-025-87719-9

Eaton, H.E., Kobayashi, T., Dermody, T.S., Johnston, R.N., Jais, P.H., Shmulevitz, M., 2017. African Swine Fever Virus NP868R Capping Enzyme Promotes Reovirus Rescue during Reverse Genetics by Promoting Reovirus Protein Expression, Virion Assembly, and RNA Incorporation into Infectious Virions. J Virol 91, e02416–16. 10.1128/JVI.02416-16

Emmott, E., De Rougemont, A., Hosmillo, M., Lu, J., Fitzmaurice, T., Haas, J., Goodfellow, I., 2019. Polyprotein processing and intermolecular interactions within the viral replication complex spatially and temporally control norovirus protease activity. Journal of Biological Chemistry 294, 4259–4271. 10.1074/jbc.RA118.006780

Emmott, E., Sweeney, T.R., Goodfellow, I., 2015. A Cell-based Fluorescence Resonance Energy Transfer (FRET) Sensor Reveals Inter- and Intragenogroup Variations in Norovirus Protease Activity and Polyprotein Cleavage. Journal of Biological Chemistry 290, 27841–27853. 10.1074/jbc.M115.688234

Goodfellow, I., Chaudhry, Y., Gioldasi, I., Gerondopoulos, A., Natoni, A., Labrie, L., Laliberté, J., Roberts, L., 2005. Calicivirus translation initiation requires an interaction between VPg and eIF4E. EMBO Reports 6, 968–972. 10.1038/sj.embor.7400510

Goodfellow, I., Taube, S., 2016. Chapter 3.2 - Calicivirus Replication and Reverse Genetics, in: Svensson, L., Desselberger, U., Greenberg, H.B., Estes, M.K. (Eds.), Viral Gastroenteritis. Academic Press, Boston, pp. 355–378. 10.1016/B978-0-12-802241-2.00017-1

Habjan, M., Penski, N., Spiegel, M., Weber, F., 2008. T7 RNA polymerase-dependent and - independent systems for cDNA-based rescue of Rift Valley fever virus. Journal of General Virology 89, 2157–2166. 10.1099/vir.0.2008/002097-0

Haga, K., Fujimoto, A., Takai-Todaka, R., Miki, M., Doan, Y.H., Murakami, K., Yokoyama, M., Murata, K., Nakanishi, A., Katayama, K., 2016. Functional receptor molecules CD300lf and CD300ld within the CD300 family enable murine noroviruses to infect cells. Proc. Natl. Acad. Sci. U.S.A. 113. 10.1073/pnas.1605575113

Hoffmann, E.S., Pascali, M.C.D., Neu, L., Domnick, C., Soldà, A., Kath-Schorr, S., 2024. Reverse transcription as key step in RNA in vitro evolution with unnatural base pairs††Electronic supplementary information (ESI) available. See DOI: https://doi.org/10.1039/d4cb00084f. RSC Chemical Biology 5, 556–566. 10.1039/d4cb00084f

Hwang, S., Alhatlani, B., Arias, A., Caddy, S.L., Christodoulou, C., Bragazzi Cunha, J., Emmott, E., Gonzalez-Hernandez, M., Kolawole, A., Lu, J., Rippinger, C., Sorgeloos, F., Thorne, L., Vashist, S., Goodfellow, I., Wobus, C.E., 2014. Murine Norovirus: Propagation, Quantification, and Genetic Manipulation. CP Microbiology 33. 10.1002/9780471729259.mc15k02s33

Kanai, Y., Komoto, S., Kawagishi, T., Nouda, R., Nagasawa, N., Onishi, M., Matsuura, Y., Taniguchi, K., Kobayashi, T., 2017. Entirely plasmid-based reverse genetics system for rotaviruses. Proc. Natl. Acad. Sci. U.S.A. 114, 2349–2354. 10.1073/pnas.1618424114

Karst, S.M., 2010. Pathogenesis of Noroviruses, Emerging RNA Viruses. Viruses 2, 748 –781. 10.3390/v2030748

Katayama, K., Murakami, K., Sharp, T.M., Guix, S., Oka, T., Takai-Todaka, R., Nakanishi, A., Crawford, S.E., Atmar, R.L., Estes, M.K., 2014. Plasmid-based human norovirus reverse genetics system produces reporter-tagged progeny virus containing infectious genomic RNA. Proc. Natl. Acad. Sci. U.S.A. 111. 10.1073/pnas.1415096111

Lee, K.H., Song, J., Kim, S., Han, S.R., Lee, S.-W., 2023. Real-time monitoring strategies for optimization of in vitro transcription and quality control of RNA. Front. Mol. Biosci. 10, 1229246. 10.3389/fmolb.2023.1229246

Lenk, R., Kleindienst, W., Szabó, G.T., Baiersdörfer, M., Boros, G., Keller, J.M., Mahiny, A.J., Vlatkovic, I., 2024. Understanding the impact of in vitro transcription byproducts and contaminants. Front. Mol. Biosci. 11, 1426129. 10.3389/fmolb.2024.1426129

Liu, G., Ni, Z., Yun, T., Yu, B., Chen, L., Zhao, W., Hua, J., Chen, J., 2008. A DNA-launched reverse genetics system for rabbit hemorrhagic disease virus reveals that the VP2 protein is not essential for virus infectivity. Journal of General Virology 89, 3080–3085. 10.1099/vir.0.2008/003525-0

Majerciak, V., Zheng, Z.-M., 2025. Induction of translation-suppressive G3BP1+ stress granules and interferon-signaling cGAS condensates by transfected plasmid DNA. hLife 3, 21–37. 10.1016/j.hlife.2024.11.005

Meier, F., Brunner, A.-D., Koch, S., Koch, H., Lubeck, M., Krause, M., Goedecke, N., Decker, J., Kosinski, T., Park, M.A., Bache, N., Hoerning, O., Cox, J., Räther, O., Mann, M., 2018. Online Parallel Accumulation–Serial Fragmentation (PASEF) with a Novel Trapped Ion Mobility Mass Spectrometer. Molecular & Cellular Proteomics 17, 2534–2545. 10.1074/mcp.TIR118.000900

Mu, X., Hur, S., 2021. Immunogenicity of *In Vitro* -Transcribed RNA. Acc. Chem. Res. 54, 4012–4023. 10.1021/acs.accounts.1c00521

Oka, T., Okemoto-Nakamura, Y., Takagi, H., 2025. CD36 is required for human sapovirus propagation. J Virol 99, e01325–25. 10.1128/jvi.01325-25

Oka, T., Stoltzfus, G.T., Zhu, C., Jung, K., Wang, Q., Saif, L.J., 2018. Attempts to grow human noroviruses, a sapovirus, and a bovine norovirus in vitro. PLoS ONE 13, e0178157. 10.1371/journal.pone.0178157

Oka, T., Takagi, H., Tohya, Y., 2014. Development of a novel single step reverse genetics system for feline calicivirus. Journal of Virological Methods 207, 178–181. 10.1016/j.jviromet.2014.07.004

Olspert, A., Hosmillo, M., Chaudhry, Y., Peil, L., Truve, E., Goodfellow, I., 2016. Protein-RNA linkage and posttranslational modifications of feline calicivirus and murine norovirus VPg proteins. PeerJ 4, e2134. 10.7717/peerj.2134

Omatola, C.A., Mshelbwala, P.P., Okolo, M.-L.O., Onoja, A.B., Abraham, J.O., Adaji, D.M., Samson, S.O., Okeme, T.O., Aminu, R.F., Akor, M.E., Ayeni, G., Muhammed, D., Akoh, P.Q., Ibrahim, D.S., Edegbo, E., Yusuf, L., Ocean, H.O., Akpala, S.N., Musa, O.A., Adamu, A.M., 2024. Noroviruses: Evolutionary Dynamics, Epidemiology, Pathogenesis, and Vaccine Advances—A Comprehensive Review. Vaccines 12, 590. 10.3390/vaccines12060590

Orchard, R.C., Wilen, C.B., Doench, J.G., Baldridge, M.T., McCune, B.T., Lee, Y.-C.J., Lee, S., Pruett-Miller, S.M., Nelson, C.A., Fremont, D.H., Virgin, H.W., 2016. Discovery of a proteinaceous cellular receptor for a norovirus. Science 353, 933–936. 10.1126/science.aaf1220

Peeters, B., De Leeuw, O., 2017. A single-plasmid reverse genetics system for the rescue of non-segmented negative-strand RNA viruses from cloned full-length cDNA. Journal of Virological Methods 248, 187–190. 10.1016/j.jviromet.2017.07.008

Peñaflor-Téllez, Y., Miguel-Rodríguez, C.E., Gutiérrez-Escolano, A.L., 2022. The Caliciviridae Family, in: Rezaei, N. (Ed.), Encyclopedia of Infection and Immunity. Elsevier, Oxford, pp. 192–206. 10.1016/B978-0-12-818731-9.00027-6

Ramanathan, A., Robb, G.B., Chan, S.-H., 2016. mRNA capping: biological functions and applications. Nucleic Acids Research 44, 7511–7526. 10.1093/nar/gkw551

Römer-Oberdörfer, A., Mundt, E., Mebatsion, T., Buchholz, U.J., Mettenleiter, T.C., 1999. Generation of recombinant lentogenic Newcastle disease virus from cDNA. Journal of General Virology 80, 2987–2995. 10.1099/0022-1317-80-11-2987

Royall, E., Locker, N., 2016. Translational Control during Calicivirus Infection. Viruses 8, 104. 10.3390/v8040104

Sandoval-Jaime, C., Green, K.Y., Sosnovtsev, S.V., 2015. Recovery of murine norovirus and feline calicivirus from plasmids encoding EMCV IRES in stable cell lines expressing T7 polymerase. Journal of Virological Methods 217, 1–7. 10.1016/j.jviromet.2015.02.003

Scribano, F.J., Gebert, J.T., Engevik, K.A., Hayes, N.M., Villanueva, J., Pham, S., Kaundal, S., Dave, J.J., Prasad, B.V.V., Estes, M.K., Ramani, S., Hyser, J.M., 2025. BTP2 restricts Tulane virus and human norovirus replication independent of store-operated calcium entry. J Virol 99, e00444–25. 10.1128/jvi.00444-25

Semenova, N., Bosnjak, M., Markelc, B., Znidar, K., Cemazar, M., Heller, L., 2019. Multiple cytosolic DNA sensors bind plasmid DNA after transfection. Nucleic Acids Research 47, 10235–10246. 10.1093/nar/gkz768

Sosnovtsev, S., Green, K.Y., 1995. RNA Transcripts Derived from a Cloned Full-Length Copy of the Feline Calicivirus Genome Do Not Require VpG for Infectivity. Virology 210, 383–390. 10.1006/viro.1995.1354

Takagi, H., Oka, T., Shimoike, T., Saito, H., Kobayashi, T., Takahashi, T., Tatsumi, C., Kataoka, M., Wang, Q., Saif, L.J., Noda, M., 2020. Human sapovirus propagation in human cell lines supplemented with bile acids. Proc. Natl. Acad. Sci. U.S.A. 117, 32078–32085. 10.1073/pnas.2007310117

Tate, J., Boldt, R.L., McFadden, B.D., D’Costa, S.M., Lewandowski, N.M., Shatzer, A.N., Gollnick, P., Condit, R.C., 2016. Biochemical analysis of the multifunctional vaccinia mRNA capping enzyme encoded by a temperature sensitive virus mutant. Virology 487, 27–40. 10.1016/j.virol.2015.10.011

Téllez, Y.P., Ibáñez, C.P., Gutiérrez-Escolano, A.L., 2024. Molecular Biology of Caliciviruses: Cellular and Viral Proteins Involved in the Establishment of the Infection, in: Pujol, F.H., Paniz-Mondolfi, A.E. (Eds.), Emerging Viruses in Latin America: Contemporary Virology. Springer Nature Switzerland, Cham, pp. 319–337. 10.1007/978-3-031-68419-7_14

UKHSA, 2026. National Norovirus and Rotavirus Report Week 43 Report Data To Week 42 Data Up To 19 October 2025.

Vinjé, J., Estes, M.K., Esteves, P., Green, K.Y., Katayama, K., Knowles, N.J., L’Homme, Y., Martella, V., Vennema, H., White, P.A., Consortium, I.R., 2019. ICTV Virus Taxonomy Profile: Caliciviridae. Journal of General Virology. 10.1099/jgv.0.001332

Ward, V.K., McCormick, C.J., Clarke, I.N., Salim, O., Wobus, C.E., Thackray, L.B., Virgin, H.W., Lambden, P.R., 2007. Recovery of infectious murine norovirus using pol II-driven expression of full-length cDNA. Proc. Natl. Acad. Sci. U.S.A. 104, 11050–11055. 10.1073/pnas.0700336104

Wei, C., Farkas, T., Sestak, K., Jiang, X., 2008. Recovery of Infectious Virus by Transfection of In Vitro-Generated RNA from Tulane Calicivirus cDNA. J Virol 82, 11429–11436. 10.1128/JVI.00696-08

Winder, N., Gohar, S., Muthana, M., 2022. Norovirus: An Overview of Virology and Preventative Measures. Viruses 14, 2811. 10.3390/v14122811

Yu, F., Haynes, S.E., Teo, G.C., Avtonomov, D.M., Polasky, D.A., Nesvizhskii, A.I., 2020. Fast Quantitative Analysis of timsTOF PASEF Data with MSFragger and IonQuant. Molecular & Cellular Proteomics 19, 1575–1585. 10.1074/mcp.TIR120.002048

Yun, T., Park, A., Hill, T.E., Pernet, O., Beaty, S.M., Juelich, T.L., Smith, J.K., Zhang, L., Wang, Y.E., Vigant, F., Gao, J., Wu, P., Lee, B., Freiberg, A.N., 2015. Efficient Reverse Genetics Reveals Genetic Determinants of Budding and Fusogenic Differences between Nipah and Hendra Viruses and Enables Real-Time Monitoring of Viral Spread in Small Animal Models of Henipavirus Infection. J Virol 89, 1242–1253. 10.1128/JVI.02583-14

Yunus, M.A., Chung, L.M.W., Chaudhry, Y., Bailey, D., Goodfellow, I., 2010. Development of an optimized RNA-based murine norovirus reverse genetics system. Journal of Virological Methods 169, 112–118. 10.1016/j.jviromet.2010.07.006

Zhang, Y., Zhang, T., Xiong, Y., Zheng, C., Li, L., 2025. Reverse Genetics-Based Methodology for Molecular Virology Study, in: Zheng, C. (Ed.), Molecular Virology: Methods and Protocols. Springer US, New York, NY, pp. 307–318. 10.1007/978-1-0716-4615-1_26

